# NAD: Noise-augmented direct sequencing of target nucleic acids by augmenting with noise and selective sampling

**DOI:** 10.1101/2023.12.14.571721

**Authors:** Hyunjin Shim

## Abstract

Next-generation sequencing necessitates a minimum quantity and concentration of DNA/RNA samples, typically achieved through amplification using the PCR technique. However, this amplification step introduces several drawbacks to biological insights, including PCR bias and the loss of epigenetic information. The advent of long-read sequencing technologies facilitates direct sequencing, with the primary constraint being the limited amount of DNA/RNA present in biological samples. Here, we present a novel method called Noise-Augmented Direct (NAD) sequencing that enables the direct sequencing of target DNA even when it falls below the minimum quantity and concentration required for long-read sequencing by augmenting with noise DNA and adaptive sampling. Adaptive sampling is an emerging technology of nanopore sequencing, allowing the enhanced sequencing of target DNA by selectively depleting noise DNA. In this study, we use the DNA standard of the Lambda phage genome as the noise DNA to augment samples containing low amounts of bacterial genomes (1 ng to 300 ng). The results with cost-effective flow cells indicate that NAD sequencing successfully detects the target DNA with an input quantity as low as 1 ng, and the bacterial genome of *Salmonella enterica* can be assembled to 30% completion at an accuracy of 98% with an input quantity of 3 ng. With high throughput flow cells, the bacterial genome of *Pseudonomas aeruginas* was assembled to near completion (99.9%) at an accuracy of 99.97% with an input quantity of 300 ng. This proof-of-concept study demonstrates the potential of NAD sequencing in enhancing the robustness of long-read sequencing with small input DNA/RNA samples with noise augmentation and adaptive sampling.

## Introduction

Long-read sequencing is revolutionizing DNA/RNA sequencing by simplifying the workflow of genomic research with the advantages of long reads [2]. This novel technology has contributed to numerous unsolved problems in the field of genomics, including the completion of the human genome through the Telomere-to-Telomere Consortium (T2T) [3] and the Human Pangenome Reference Consortium (HPRC) [4–6]. Among the long-read sequencing technologies, nanopore sequencing has several unique advantages by utilizing the biophysical properties of nanopores and nucleic acids [7,8]. This technology measures changes in ionic current flows across a protein nanopore as a string of nucleic acids passes through to reconstruct the genetic sequence from the electric signals [9]. This signal conversion process further employs advanced computational methods such as neural networks and parallel computing, leveraging recent technological advancements spanning various fields [10,11]. Notable advantages of nanopore sequencing over other sequencing methods include the ability to sequence native DNA/RNA and the ability to sequence selectively through depletion or enrichment. Direct sequencing of native DNA/RNA generates additional information on the sequences. This epigenetic information, such as the detection of DNA and RNA methylation, plays a key role in understanding many diseases, including cancer [12,13]. Furthermore, nanopore sequencing has an important feature of adaptive sampling, which selectively depletes or enriches sequences of interest [14]. This feature relies on the ability to control each pore by reversing the current across it and to process basecalling and mapping extremely fast with neural networks and parallel computing [10].

Despite these advantages, nanopore sequencing also requires a high input quantity of DNA/RNA to fully leverage these features of direct sequencing and adaptive sampling. For instance, most protocols recommend an input DNA/RNA quantity of at least 500 ng and 1,000 ng for Flongle flow cells and MinION flow cells, respectively. This high input quantity of DNA/RNA is necessary for the adapter efficiency in the ligation preparation, as well as the long-read sequencing of DNA/RNA [15]. This requirement is a limiting factor in utilizing nanopore sequencing in genomic studies involving biological samples with small quantities of DNA/RNA. In scenarios involving small DNA/RNA quantities within samples, an extra PCR step is often employed to amplify these scarce sequences, undermining the capacity of nanopores to capture epigenetic information. Moreover, it necessitates the quantification of DNA/RNA during each sequencing experiment, typically with a stringent quantity threshold. These experimental criteria often lead to the abandonment of direct sequencing or long-read sequencing in numerous studies [16].

Here, we introduce a novel method called Noise-Augmented Direct (NAD) sequencing, which augments a sample of a small DNA quantity (<500 ng) with the ‘noise’ DNA to selectively sequence only the target sequence using adaptive sampling (Figure 1). The noise or background DNA, such as human DNA during sequencing experiments, is commonly considered an undesirable form of contamination that needs to be identified and eliminated [17,18]. However, the concept of adding noise to increase the generalization performance with real-world data is widely utilized in other fields. For instance, data augmentation with white noise is known to improve the accuracy of deep-learning models in real-world testing [19]. Deep learning has been widely used for various artificial intelligence (AI) tasks such as speech and image recognition and natural language processing [20]. However, the generalization performance of deep learning reduces drastically when tested in noisy data, which is often the case in real-world data [21]. One of the strategies to circumvent this pitfall is to train the neural networks using data augmented with some random noise to increase the robustness and generalization power [22].

**Figure 1:**
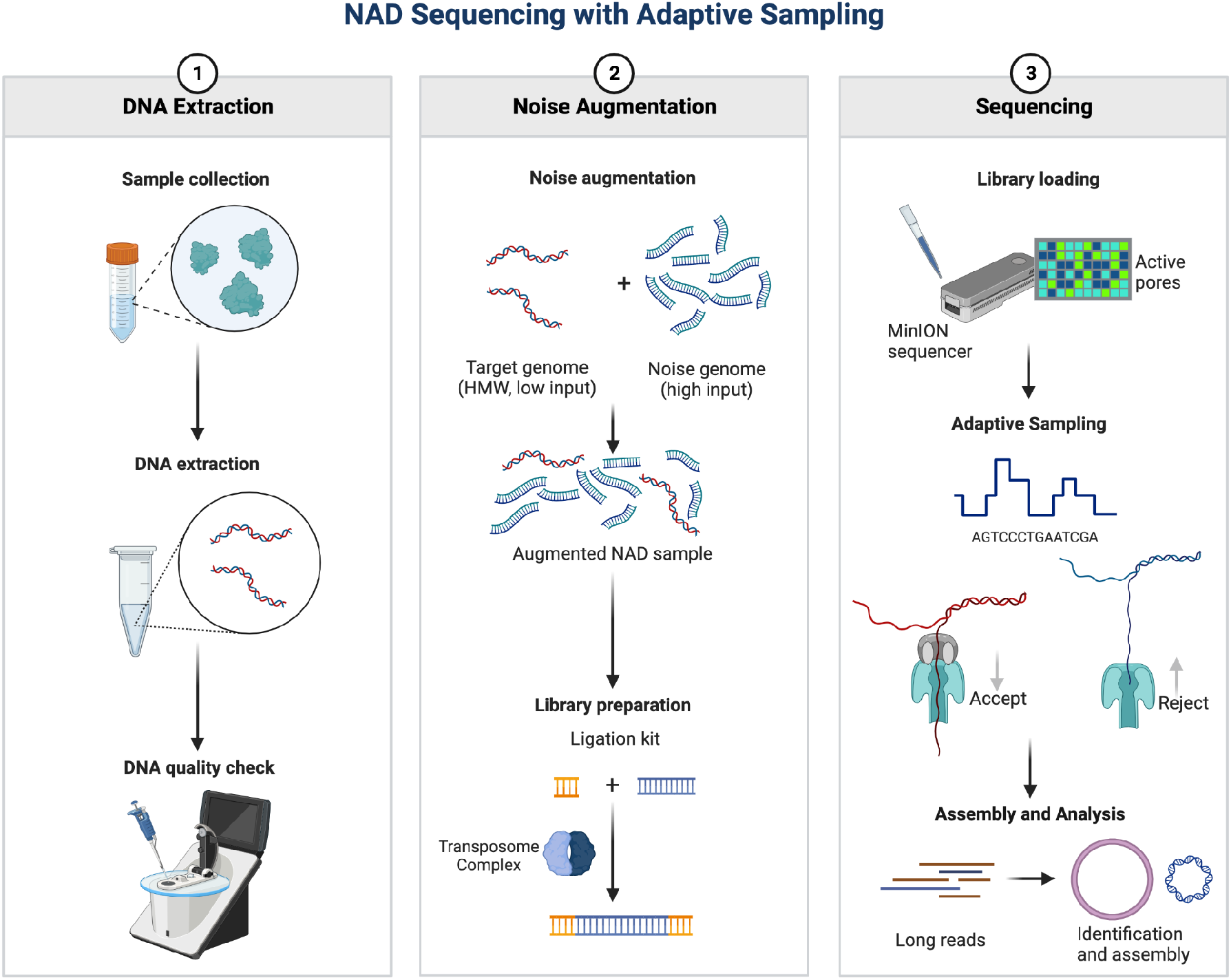
Process of Noise-Augmented Direct (NAD) sequencing. The target DNA/RNA is extracted from a sample of interest (e.g. human nasal swab) and augmented with the noise DNA/RNA. The sequencing library of the target and noise DNA/RNA is directly sequenced using adaptive sampling by depleting the noise DNA/RNA. This allows direct sequencing of the target DNA/RNA in a native form, without the need to amplify the target DNA/RNA that is below the minimum quantity and concentration of long-read sequencing requirements.

In this study, we adopt a similar line of reasoning to increase the generalization performance of nanopore sequencing in real-world datasets by adding a controlled amount of ‘noise’ DNA to a biological sample and exploiting the ability of adaptive sampling to enrich the target DNA by selectively depleting the noise DNA. We use the lambda phage genome as the noise DNA and two bacterial genomes (*Pseudomonas aeruginosa* and *Salmonella enterica*) as the target DNA. We demonstrate that NAD sequencing can detect the target DNA with an input quantity as low as 1 ng in cost-effective Flongle flow cells. Furthermore, NAD sequencing can efficiently assemble parts of the bacterial genome, with a target DNA quantity as low as 3 ng (30% complete with an accuracy of 97.58%) in cost-effective Flongle flow cells, and a target DNA quantity of 300 ng (99.9%% complete with an accuracy of 99.97%) in high-throughput MinION flow cells. This result demonstrates the potential of NAD sequencing as a practical method for detecting and assembling target DNA of limited quantity. This proof-of-concept study underscores the effectiveness of injecting noise DNA to augment target DNA to extend the applicability of long-read sequencing to real-world settings where biological samples often contain limited nucleic acids. We further discuss the necessity of enhancing the integration of computational processing power to handle the vast datasets generated by NAD sequencing, particularly when incorporating adaptive sampling in real-time.

## Results

### Quality control of NAD sequencing experiments

Microbial DNA standards serve as a standard for benchmarking the performance along the workflow of genomics analyses and as a tool to increase reproducibility. Before noise augmentation, the quality and quantity of the DNA standards were checked using a DNA Spectrophotometer. The results show that the DNA standard of the noise genome (Lambda phage) was as stated by the manufacturers at the concentration of ∼600 ng/μL, meeting the quality criteria of 260/280 and 260/230 at ∼1.8 and ∼2.0, respectively (Figure S1 and Table S1). However, the target DNA samples were below the minimum concentration range of the DNA Spectrophotometer. For example, the target DNA sample of *P. aeruginosa* was measured to be 15.322 ng/μL before augmenting with the noise DNA (Table S1), which was close to the concentration stated by the manufacturers of 10 ng/μL. Due to the minimum concentration threshold, the quality control values of 260/280 and 260/230 were not relevant for these target DNA samples.

### The data output of NAD sequencing experiments

The target DNA samples that were below the minimum concentration were augmented with the noise DNA to meet the input DNA criteria for nanopore sequencing (Table 1). The augmented DNA samples of *Pseudomonas aeruginosa* and *Salmonella enterica* were ligated and sequenced in MinION flow cells and Flongle flow cells, respectively.

**Table 1:**
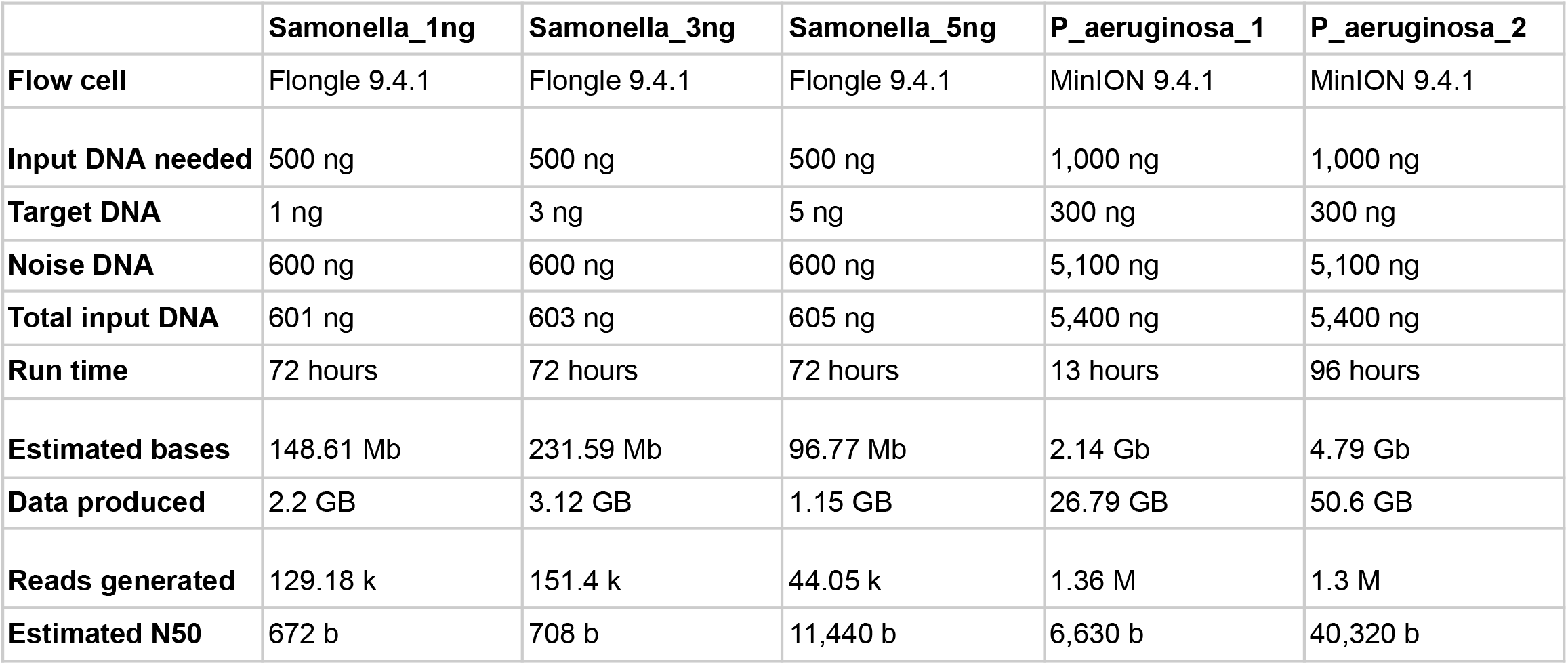
Experimental parameters of NAD sequencing experiments.

|  | **Samonella_1ng** | **Samonella_3ng** | **Samonella_5ng** | **P_aeruginosa_1** | **P_aeruginosa_2** |
| --- | --- | --- | --- | --- | --- |
| **Flow cell** | Flongle 9.4.1 | Flongle 9.4.1 | Flongle 9.4.1 | MinION 9.4.1 | MinION 9.4.1 |
| **Input DNA needed** | 500 ng | 500 ng | 500 ng | 1,000 ng | 1,000 ng |
| **Target DNA** | 1 ng | 3 ng | 5 ng | 300 ng | 300 ng |
| **Noise DNA** | 600 ng | 600 ng | 600 ng | 5,100 ng | 5,100 ng |
| **Total input DNA** | 601 ng | 603 ng | 605 ng | 5,400 ng | 5,400 ng |
| **Run time** | 72 hours | 72 hours | 72 hours | 13 hours | 96 hours |
| **Estimated bases** | 148.61 Mb | 231.59 Mb | 96.77 Mb | 2.14 Gb | 4.79 Gb |
| **Data produced** | 2.2 GB | 3.12 GB | 1.15 GB | 26.79 GB | 50.6 GB |
| **Reads generated** | 129.18 k | 151.4 k | 44.05 k | 1.36 M | 1.3 M |
| **Estimated N50** | 672 b | 708 b | 11,440 b | 6,630 b | 40,320 b |

The noise-augmented sample of *P. aeruginosa* had two technical replicates, where the first replicate was only run for 13 hours due to an error in flow cells (Table 1). The second replicate was run for 96 hours to completely exploit the capacity of the MinION flow cell. However, the quality score of real-time basecalling declined significantly after ∼72 hours despite replenishing with the flush buffer, which is a typical runtime of MinION flow cells (data not shown). The second replicate has a better experimental output with twice more bases sequenced (4.79 Gb) as compared to the first replicate (2.14 Gb). This data output was below the theoretical output of MinION flow cells at 50 Gb, but achieving 10% of the theoretical capacity was typical data sizes generated with nanopore sequencing in the previous experiments [16]. The second replicate also has a better N50 estimate (40,320 b) which was around 5 times longer than that of the first replicate (Figure S2). The estimated N50 from the reads shows a satisfactory performance of high-molecular-weight DNA extraction and sequencing.

The noise-augmented sample of *S. enterica* had three experimental runs with a range of target DNA concentrations (1 ng, 3 ng, and 5 ng). All the experiments were run for 72 hours to completely exploit the capacity of the Flongle flow cells. However, the quality score of real-time basecalling declined significantly after ∼24 hours, which is a typical runtime of Flongle flow cells (data not shown). All the runs generated around 5% to 10% of the theoretical capacity of Flongle flow cells at 2.8 Gb (Table 1). The N50 estimates of all the sequencing runs showed much lower read lengths than expected of high-molecular-weight DNAs (Figure S2). The sequencing run with the largest amount of target DNA at 5ng shows the largest estimated N50 at 11,440 b. The estimated N50 correlates with the concentration of the target DNA, indicating that the low estimated N50 may arise as an artifact of adaptive sampling of the noise DNA of the Lambda phage genome.

### Adaptive sampling of NAD sequencing experiments

In NAD sequencing of these noise-augmented samples, the adaptive sampling feature of nanopore sequencers was used to enrich the target DNA by depleting the noise DNA. The read length graphs from the nanopore report show there are two peaks in all of the NAD sequencing experiments (Figure S2). The first peaks are at the read length of 500 b, while the second peaks are at the read length of ∼45 kb. The bimodal distribution of these read length graphs indicates successful adaptive sampling in these experiments, with the first peak representing the noise DNA that was rejected within a few seconds after being sequenced with a nanopore. The second peak likely represents the target DNA of bacterial genomes from the DNA standards.

During adaptive sampling, each read passing the nanopore was mapped against the reference genome of the Lambda phage while sequencing (Table S4). There are three types of decisions in adaptive sampling: “no_decision” when the read has been continued and mapped against the reference(s), “stop_receiving” when the read was accepted and fully sequenced, and “unblock” when the read was stopped and rejected by reversing of the voltage.

To evaluate the decision-making process of adaptive sampling, the proportion of these three types of decisions was plotted for each NAD sample, in nominal value (Figure 2A and Figure S3) and relative value (Figure 2B). For all the samples, a majority of the reads were “unblocked” while being sequenced, indicating that more than 50% of reads were rejected as adaptive sampling is set to deplete the noise DNA. For the Salmonella samples, there is a decreasing trend of unblocked reads as the concentration of the target DNA increases. This trend indicates that adaptive sampling was functioning as expected, as the nanopores rejected a fewer number of reads when there were fewer noise DNAs present in the sample. Conversely, the proportion of acceptance (“stop_receiving”) in sequencing increases as the concentration of the target DNA increases in the Salmonella samples. For NAD sequencing, it is notable that the proportion of unblocked reads is much larger than the proportion of accepted reads, as adaptive sampling is set to deplete rather than enrich.

**Figure 2:**
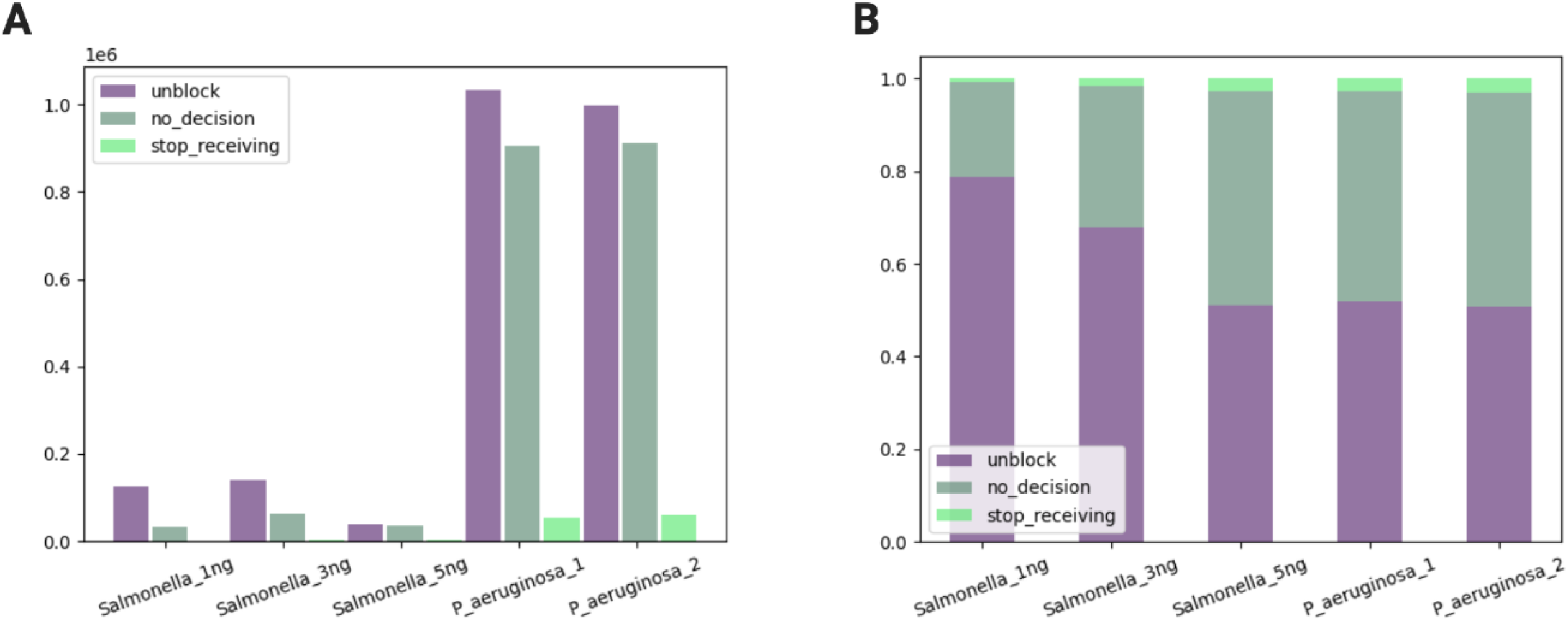
Three decision types in adaptive sampling of the NAD samples. (A) comparison in nominal value; (B) comparison in relative value. “no_decision” when the read has been continued and mapped against the reference(s); “stop_receiving” when the read was accepted and fully sequenced; “unblock” when the read was stopped and rejected by the nanopore.

### Performance of NAD sequencing experiments

To evaluate the performance of NAD sequencing, the sequence length of the reads was compared by the sample type (Figure 3A) and by the classification type (Figure 3B). In all of the samples, the average sequence length of the reads that were unblocked and rejected as the noise DNA was much shorter at 500 b than those of the other reads (Figure 3A). The nanopore sequencer was preset to deplete the noise DNA by rejecting the Lambda phage genome of ∼50 kb using adaptive sampling. As it takes around 1 second for nanopores to decide on whether to accept or reject the read using adaptive sampling, this result verifies the implementation of adaptive sampling given the translocation speed of 450 bases per second [23]. Furthermore, the average sequence length of the reads that were accepted and fully sequenced (“stop_receiving”) was much higher at 4,000 b than that of the reads that were continued (“no_decision”) at 1,000 b. This shows that the reads that were actively accepted in adaptive sampling are more likely to be the target DNA of bacterial genomes than the reads that were passively continued, potentially resulting from the bottleneck of basecalling for read mapping.

**Figure 3:**
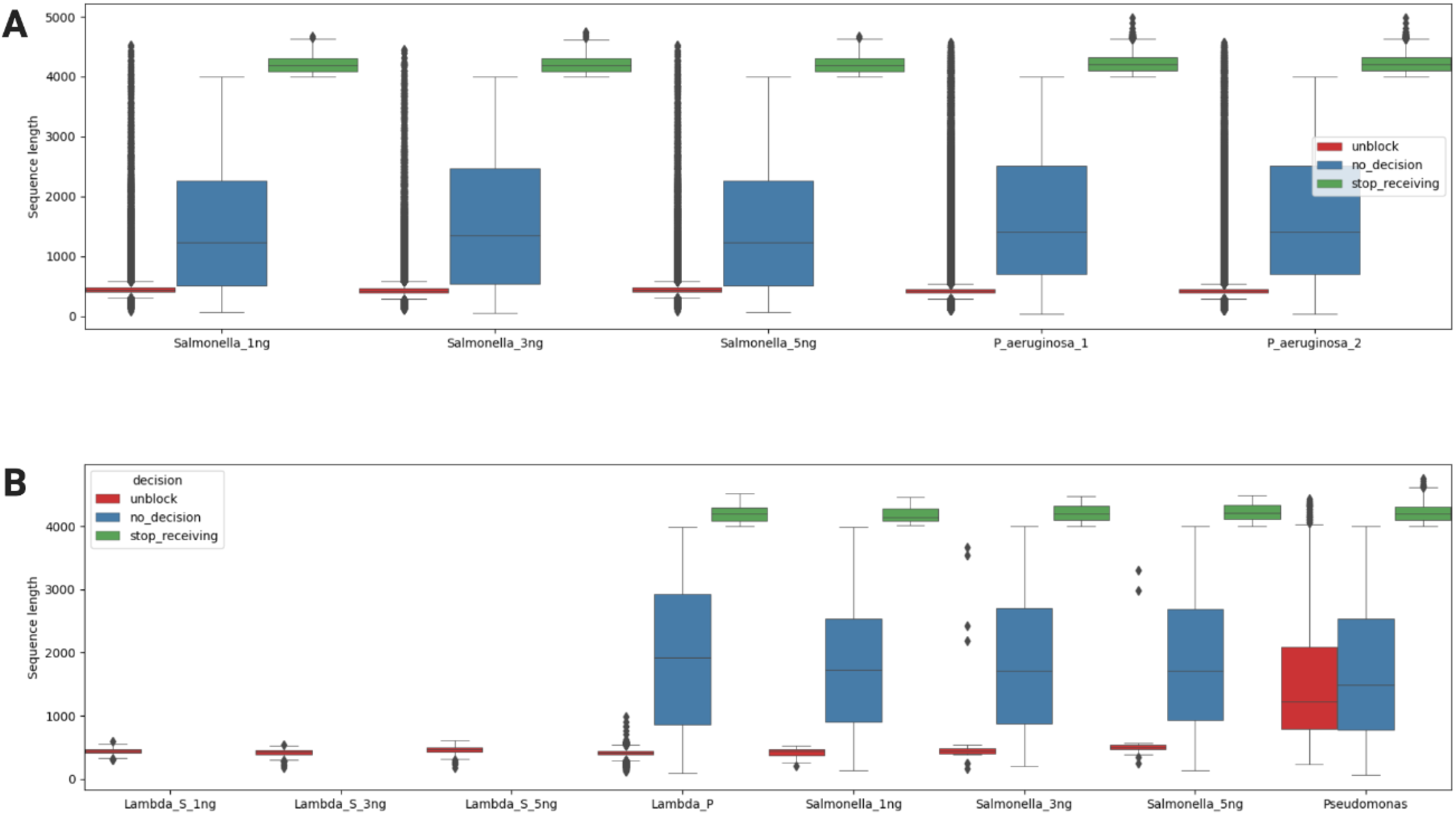
Sequence length of the NAD sequencing experiments. (A) Comparison by the sample: (B) Comparison by the noise DNA (Lambda phage) versus the target DNA (*S. enterica* or *P. aeruginosa*). Lambda_S indicates the noise DNA from the samples of *S. enterica* and Lambda_P indicates the noise DNA from the sample of *P. aeruginosa*

To confirm the decision-making process of adaptive sampling, each read was classified by a cloud-based analysis platform called the WIMP workflow (Figures S4 and S5). The species identification was downloaded from the WIMP workflow, and the reads identified as the noise DNA or the target DNA were saved separately. Subsequently, each identified read was subcategorized into its adaptive sampling decision type (Figure 3B). The results show that the noise DNA from the Salmonella samples was all rejected (“unblocked”) by adaptive sampling. Interestingly, there were some noise DNA reads from the Pseudomonas sample that were either continued (“no_decision”) or accepted (“stop_receiving”). This result is unexpected as MinION flow cells are expected to have more stable nanopores and generate higher outputs. However, the current setting of GPU-accelerated adaptive sampling may be a limiting factor in depleting the noise DNA with a larger number of nanopores generating big data output in NAD sequencing experiments.

For the target DNA, the NAD experiments successfully continued (“no_decision”) or accepted (“stop_receiving”) a larger number of reads in both the Salmonella samples and the Pseudomonas sample (Table S5). In the Pseudomonas sample, the target DNA reads with longer sequence lengths were rejected wrongly (“unblocked”) than those of the Salmonella samples (Figure 3B), which indicates that the decision-making process of adaptive sampling takes longer in higher throughput MinION flow cells than Flongle flow cells, due to the limitation of parallel computing power in this study.

In the Salmonella samples, the output ratio of the target DNA to the noise DNA increases from 2.69 to 33.17 as the concentration of the target DNA increases from 1 ng to 5 ng (Table 2). Furthermore, the ratio of acceptance to rejection of the target DNA increases from 1.30 to 20.58 with the increasing concentration of the bacterial DNA content. This result indicates that adaptive sampling functions with predictable outcomes, and NAD experiments perform effectively in sequencing low concentrations of target DNAs. Additionally, the output ratio of the target DNA length and the noise DNA length remain constant at around 4 (Table 2), indicating this correlation in the performance and the concentration of the target DNA is independent of other factors in NAD experiments.

**Table 2:**
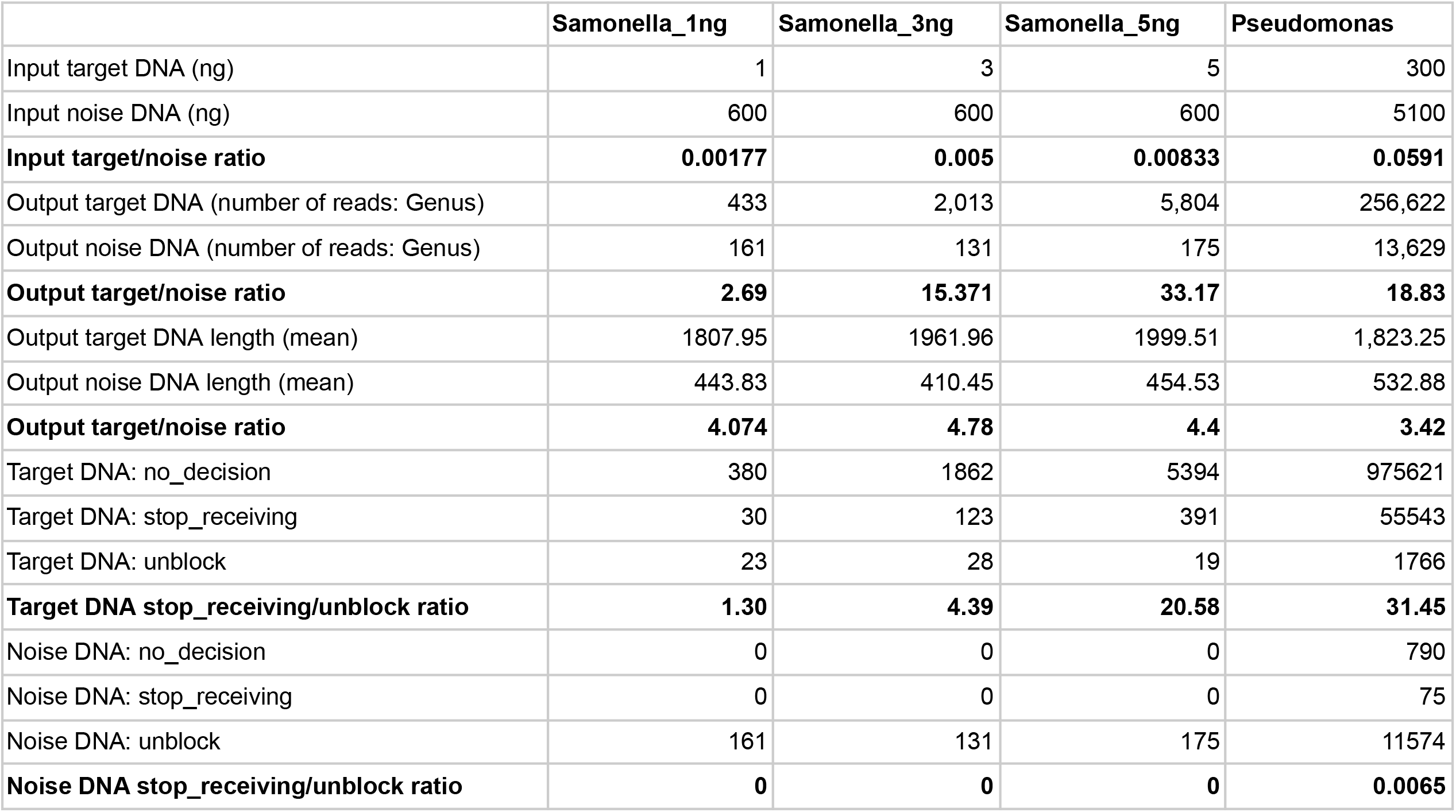
Summary statistics of the NAD datasets by WIMP classification at the Genus level (Target DNA: *Salmonella enterica* or *Pseudomonas aeruginosa,* Noise DNA: Lambdavirus)

|  | Samonella_1ng | Samonella_3ng | Samonella_5ng | Pseudomonas |
| --- | --- | --- | --- | --- |
| Input target DNA (ng) | 1 | 3 | 5 | 300 |
| Input noise DNA (ng) | 600 | 600 | 600 | 5100 |
| **Input target/noise ratio** | **0.00177** | **0.005** | **0.00833** | **0.0591** |
| Output target DNA (number of reads: Genus) | 433 | 2,013 | 5,804 | 256,622 |
| Output noise DNA (number of reads: Genus) | 161 | 131 | 175 | 13,629 |
| **Output target/noise ratio** | **2.69** | **15.371** | **33.17** | **18.83** |
| Output target DNA length (mean) | 1807.95 | 1961.96 | 1999.51 | 1,823.25 |
| Output noise DNA length (mean) | 443.83 | 410.45 | 454.53 | 532.88 |
| **Output target/noise ratio** | **4.074** | **4.78** | **4.4** | **3.42** |
| Target DNA: no_decision | 380 | 1862 | 5394 | 975621 |
| Target DNA: stop_receiving | 30 | 123 | 391 | 55543 |
| Target DNA: unblock | 23 | 28 | 19 | 1766 |
| **Target DNA stop_receiving/unblock ratio** | **1.30** | **4.39** | **20.58** | **31.45** |
| Noise DNA: no_decision | 0 | 0 | 0 | 790 |
| Noise DNA: stop_receiving | 0 | 0 | 0 | 75 |
| Noise DNA: unblock | 161 | 131 | 175 | 11574 |
| **Noise DNA stop_receiving/unblock ratio** | **0** | **0** | **0** | **0.0065** |

However, the Pseudomonas sample has a much lower output ratio of the target DNA to the noise DNA at 18.83 despite a higher input ratio of the target DNA to the noise DNA. This result is due to the noise DNA being accepted or fully sequenced, as shown by the higher ratio of acceptance to rejection in the noise DNA (Tables 2 and S5). These summary statistics emphasize the importance of increasing parallel computing power to process higher throughput in MinION flow cells for NAD experiments.

### Metagenomic assembly of the target DNA and the noise DNA from NAD sequencing experiments

The species identification of each read using the WIMP workflow shows that while a large number of reads was correctly classified as the target DNA or the noise DNA at the family level, a higher proportion of reads was wrongly classified as other organisms at the genus level (Figures S4 and S5). For example, a majority of the reads were identified as *Escherichia coli* in the Salmonella samples. This misclassification may arise from the fact the genomes of *Escherichia coli* and *Salmonella enterica* share similar gene content [24,25], making it difficult for the Centrifuge classification engine to accurately identify the genus or species using only a single long read.

After the rapid species classification, the potential of genome assembly using NAD sequencing experiments with such low DNA inputs was investigated. The reads from the NAD datasets were assembled with a de novo assembler for single-molecule sequencing reads called Flye using a metagenomic option. The assembled fragments had an N50 of around 50 kb in the Salmonella samples and an N50 of around 1 Mb in the Pseudomoa sample (Table S6). This shows that MinION flow cells are much more effective in assembling bacterial genomes with a higher data output, despite the lower efficiency in adaptive sampling (Table 2).

After metagenomic genome assembly, the assembled fragments were aligned against the reference target genome or the reference noise genome (Tables 3 and S7). The results show that the Salmonella samples assembled only a small percentage of the target genome of *S. enterica*. For example, the assembled fragments from the Salmonella_1ng sample only covered 0.36% of the reference target genome, while it covered 100% of the reference noise genome (Figure 4). The coverage of the target genome increases with the increasing target DNA concentrations, as shown by the Salmonella_3ng sample and the Salmonella_5ng sample covering almost 30% and 20% of the reference target genome, respectively. They both covered the reference noise genome fully as an artifact of NAD sequencing experiments (Table S8). Notably, the Pseudomonas sample covered 99.9% of the reference target genome (Figure 5).

**Figure 4:**
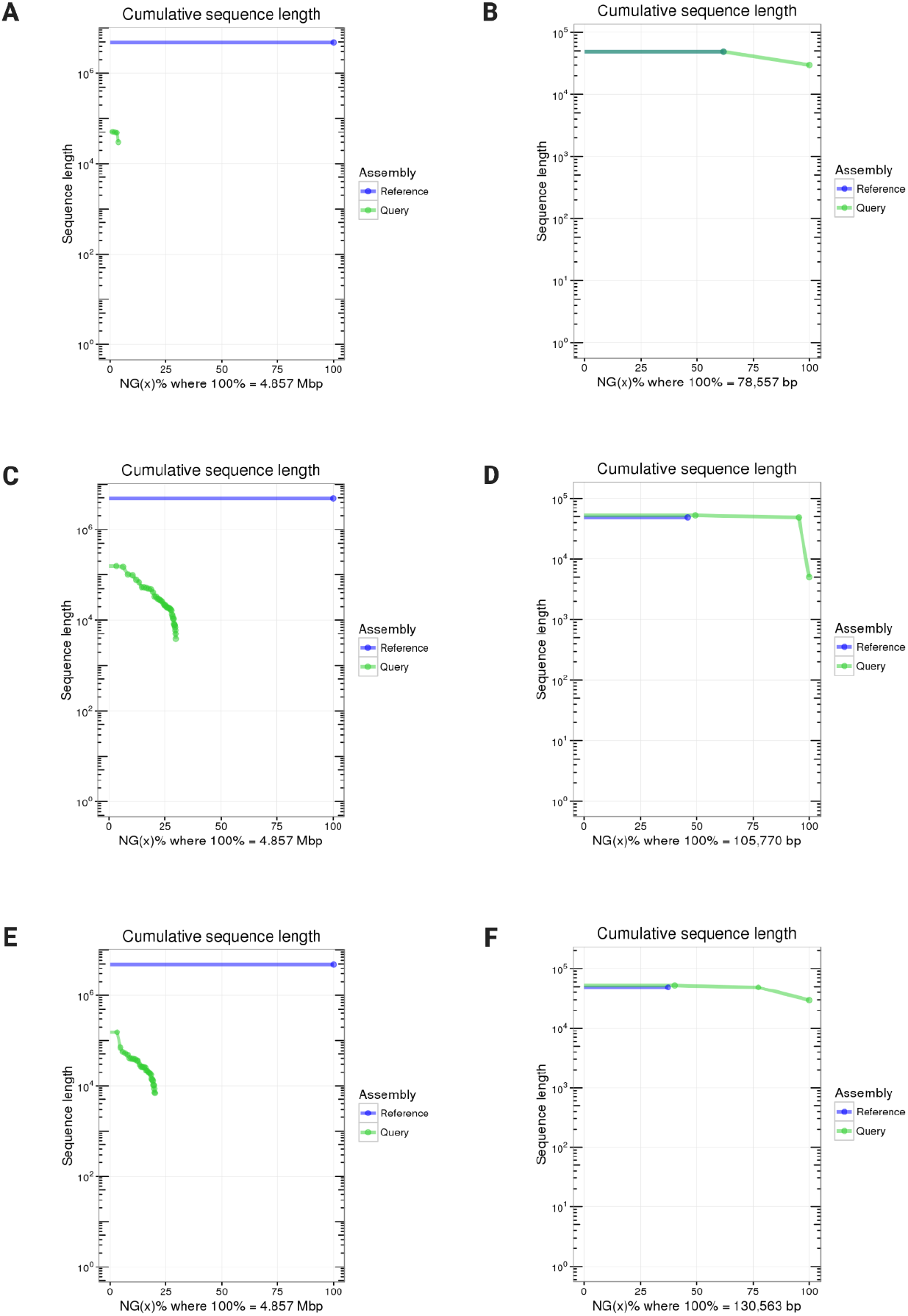
MUMmer N charts to compare Flye-assembled genomes against the target reference (*Salmonella enterica*: GCF_000006945) or the noise reference (Lambda phage: NC_001416). (A) Salmonella_1ng against the target reference; (B) Salmonella_1ng against the noise reference; (C) Salmonella_3ng against the target reference; (D) Salmonella_3ng against the noise reference; (E) Salmonella_5ng against the target reference; (F) Salmonella_5ng against the noise reference.

**Figure 5:**
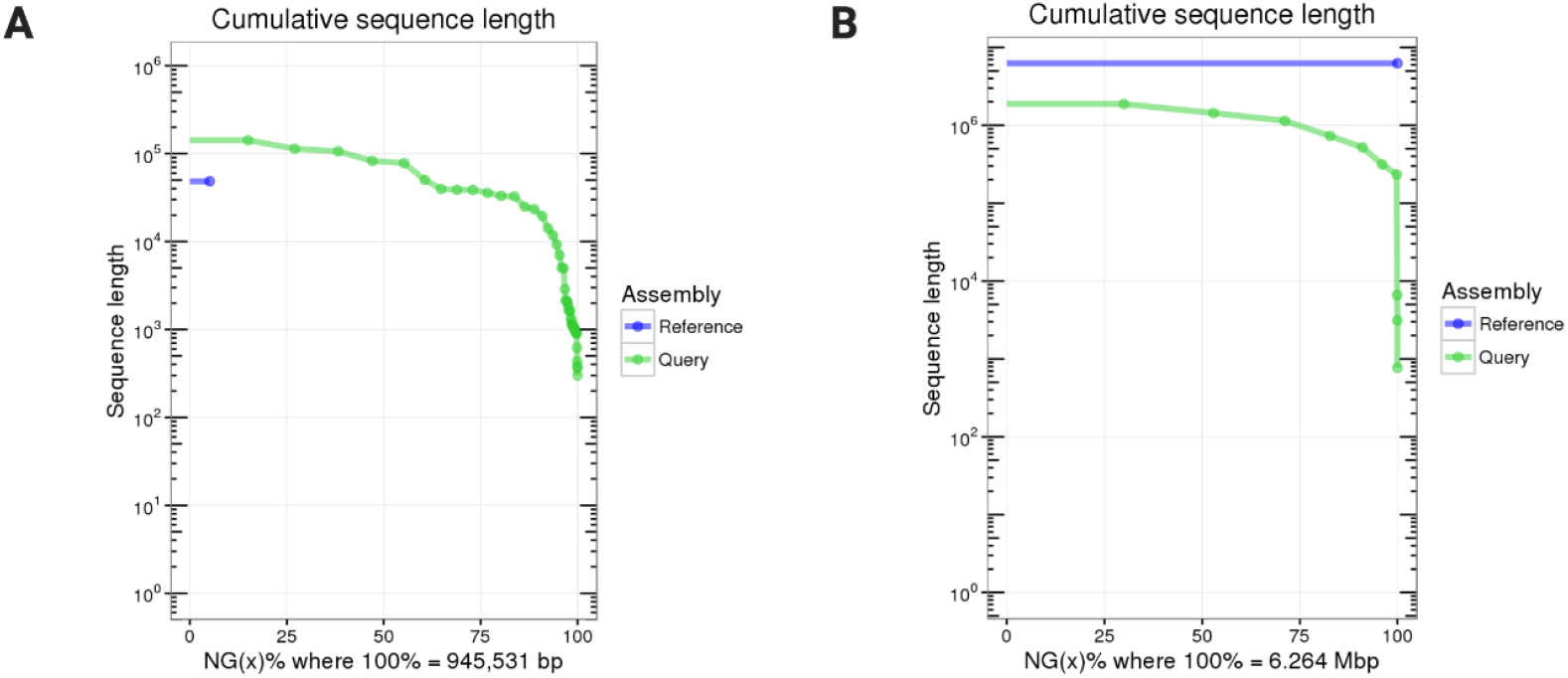
MUMmer N charts to compare Flye-assembled genomes against the target reference (*Pseudomonas aeruginosa*: GCF_000006765) or the noise reference (Lambda phage: NC_001416). (A) Pseudomonas_300 ng against the target reference; (B) Pseudomonas_300ng against the noise reference.

The assembled genome had a variable average identity depending on the sample type. The Pseudomonas sample had the highest accuracy of 99.97% in genome assembly when aligned to the reference target genome (Table 3). The assembled genome of the noise DNA also had the highest accuracy of 99.93% when mapped to the reference noise genome (Table S7). The Salmonella_1ng of the lowest target concentration had the lowest accuracy of 82.53% in genome assembly when aligned to the reference target genome (Table 3). In this NAD sample, several mutations were detected, including breakpoints, insertions, and single nucleotide polymorphisms (SNPs) (Figure S6). However, the assembled genome of the noise DNA had a high accuracy of 99.89% when mapped to the reference noise genome (Table S8). The Salmonella_3ng and the Salmonella_5ng samples had an average identity of 97.58% and 97.77% when aligned to the reference target genome, respectively (Table 3). These results show that NAD sequencing experiments accurately assemble full bacterial genomes at lower input DNAs (300 ng) than recommended (1,000 ng) using MinION flow cells. Furthermore, NAD sequencing experiments can assemble a fraction of bacterial genomes (∼30%) accurately from an incredibly low input DNAs of 3ng than recommended (500 ng) using Flongle flow cells. Lower input DNAs may still be used for species identification with NAD sequencing, potentially solving the current limitation of long-read sequencing due to the high input DNA requirement.

**Table 3:** Analysis of the assembled target genomes against the reference target genome.

|  | *S. enterica*<br>reference | Samonella_1<br>ng | *S. enterica*<br>reference | Samonella_3<br>ng | *S. enterica*<br>reference | Samonella_5<br>ng | *P. Aeruginosa*<br>reference | P_aeruginosa |
| --- | --- | --- | --- | --- | --- | --- | --- | --- |
| **Total Bases** | 4857450 | 182527 | 4857450 | 1557907 | 4857450 | 1055836 | 6264404 | 7439804 |
| **Aligned Bases** | **17573(0.36%)** | **16793(9.20%)** | **1357232(27.94%)** | **1279936(82.16%)** | **899445(18.52%)** | **846000(80.13%)** | **6258121(99.90%)** | **6297543(84.65%)** |
| **Unaligned Bases** | 4839877(99.64%) | 165734(90.80%) | 3500218(72.06%) | 277971(17.84%) | 3958005(81.48%) | 209836(19.87%) | 6283(0.10%) | 1142261(15.35%) |
| **Total Sequences** | 1 | 5 | 1 | 44 | 1 | 35 | 1 | 98 |
| **Aligned Sequences** | 1(100.00%) | 4(80.00%) | 1(100.00%) | 42(95.45%) | 1(100.00%) | 33(94.29%) | 1(100.00%) | 20(20.41%) |
| **Unaligned Sequences** | 0(0.00%) | 1(20.00%) | 0(0.00%) | 2(4.55%) | 0(0.00%) | 2(5.71%) | 0(0.00%) | 78(79.59%) |
| **1-to-1** | 10 | 10 | 59 | 59 | 48 | 48 | 14 | 14 |
| **Total Length** | 16798 | 16793 | 1280926 | 1285001 | 845297 | 845479 | 6263146 | 6261391 |
| **Average Length** | 1679.8 | 1679.3 | 21710.61 | 21779.68 | 17610.35 | 17614.15 | 447367.57 | 447242.21 |
| **Average Identity** | **82.53** | **82.53** | **97.58** | **97.58** | **97.77** | **97.77** | **99.97** | **99.97** |

## Discussion

NAD sequencing explores the potential of sequencing low-input target DNA/RNA in its native state by augmenting biological samples with a controlled quantity and quality of noise DNA/RNA. This concept is inspired by the data augmentation technique of machine learning and enabled by the technological advances in long-read sequencing and parallel computing. This study has the specific aim of lowering the minimum input DNA/RNA to a fraction of the recommended input amount of 500 ng and 1,000 ng in cost-effective and high-throughput nanopore sequencing, respectively. This input quantity requirement of DNA/RNA is not realistic in many biological samples without amplification. Conventionally, these biological samples with scarce DNA/RNA have been amplified with the Polymerase Chain Reaction (PCR) technique to meet the minimum input criteria of input DNA/RNA before sequencing.

PCR is a widely used molecular biology technique that allows the amplification of specific DNA sequences. Using PCR, very small amounts of DNA sequences are exponentially amplified to millions to billions of copies with a DNA polymerase in a series of cycles of temperature changes. PCR is fundamental to a broad variety of biotechnological research, including biomedical research, genetic testing, and forensic science [1,26]. However, there are several PCR-induced biases and artifacts, such as DNA polymerase errors and the loss of epigenetic signatures [27–29]. These PCR-induced issues have been a limiting factor in understanding some biological processes, such as DNA methylation which plays a key role in development and gene expression [30]. Recently, there has been an increasing interest in studying the genomes of various species and individuals at the epigenetic level, of how DNA and RNA sequences undergo epigenetic modifications to inherit information without changing the genetic sequences [31]. Such epigenetic information is relevant in medical fields such as cancer genomics [32,33], but recent findings also suggest that various organisms utilize base modifications to escape host immunity [34,35]. The development of the ground-breaking mRNA vaccine also arises from the differentially modified nucleotides as a method to transport mRNA without triggering the immune system [36,37]. Thus, expanding the capacity to sequence native DNA/RNA from diverse biological samples will enable the scientific community to further explore novel territories of epigenetics.

The study aims to develop a method that broadens the possibility of direct sequencing for various biological samples so that more DNA/RNA can be sequenced in their native states. In this study, the novel method of NAD sequencing tested the minimum input ranges of target DNA from 1 ng to 5 ng for cost-effective nanopore sequencing and 300 ng for high-throughput nanopore sequencing. We demonstrate that NAD sequencing can detect the target DNA with a quantity as low as 1 ng with the cost-effective Flongle flow cells. Furthermore, NAD sequencing can efficiently assemble parts of a bacterial genome, with the target DNA quantity as small as 3 ng (30% complete with an accuracy of 97.58%) in the cost-effective Flongle flow cell, and the target DNA quantity of 300 ng (99.9% complete with an accuracy of 99.97%) in the high-throughput MinION flow cell.

The initial concentration and quantity of the microbial DNA standards of *Salmonella enterica* and *Pseudomonas aeruginosa*, approximately at ∼10 ng/μl, fell below the minimum concentration measurable with the confidence level of a DNA spectrophotometer. Without NAD sequencing, the target DNA within all of these noise-augmented samples was not sufficient to be sequenced effectively in their native form with nanopore sequencing. The smallest quantity of the target DNA examined in this study was 1 ng of *Salmonella enterica* genome, constrained by the limitations of the laboratory equipment, but NAD sequencing exhibits the capability to detect smaller input amounts of the target DNA than those employed in this study.

The broad aim of this study is to increase the robustness of long-read sequencing to real-world biological samples of small input amounts and noisy backgrounds. Because of the high input requirement, the option of direct sequencing using long-read technologies is frequently supplanted by PCR and short-read sequencing. Conversely, NAD sequencing capitalizes on noise combined with adaptive sampling to mitigate the challenge associated with the high DNA inputs required in long-read sequencing, without the need for amplification or any supplementary processing. The only additional step is to determine the type and amount of noise DNA necessary to attain the noise-augmented sample requisite for adaptive sampling. Augmenting biological samples with a controlled noise DNA is inspired by the data augmentation technique of white noise injection in machine learning to improve the robustness and generalization power of deep learning models with noisy real-world data.

In future developments, the proof-of-concept of NAD sequencing experiments will be broadened to establish standardized protocols for augmenting biological samples with the noise DNA. These protocols aim to make the quantification of input DNA/RNA redundant for experimental conditions where such equipment is not available, as in cost-constraint settings or remote sampling campaigns. Nanopore sequencing has been optimized to perform in remote and cost-constraint situations where rapid and on-site sequencing of biological samples is needed. Such scenarios may entail the urgency of swiftly detecting infectious agents in remote areas. In such cases, the simple addition of the noise DNA such as the Lambda phage genome will suffice in detecting scarce target DNA/RNAs or even assembling the target genome. This noise augmentation ensures the robustness of direct sequencing in real-world biological data, as well as eliminating the bottleneck of DNA/RNA quantification.

## Methods

### Quality control of the microbial DNA standards

Microbial DNA standards from *Pseudomonas aeruginosa* and *Salmonella enterica* (Sigma-Aldrich) were obtained at the concentration of 10 ng/μL. For both the microbial DNA standards, the UV absorbance ratio (OD260/OD280) and the bacteria identity from the manufacturers are given as 1.8 and 95%, respectively. DNA concentration and purity were confirmed (Table S1, Figure S1) using a DNA spectrophotometer (DS-11 Series from DeNovix). Given the total volume of 30 μl, the total amount of DNA from these microbial standards was 300 ng. The input DNA requirements for MinION flow cells (R9.4.1) and Flongle flow cells (R9.4.1) are 1,000 ng and 500 ng, respectively.

### Noise augmentation of the microbial DNA standards

Lamda DNA standards from *Escherichia coli* bacteriophage (Thermo Fisher) were obtained at the concentration of 0.3 μg/μL. To meet the input DNA requirements of nanopore sequencing, the microbial DNA standards were augmented with the lambda DNA standard. For nanopore sequencing of *P. aeruginosa*, 30 μl of the microbial DNA standard was augmented with 17 μl of the lambda DNA standard to obtain 5.4 μg of DNA (Table S2). The final concentration of the noise-augmented sample far exceeds the minimum DNA input requirement for MinION flow cells (R9.4.1). For nanopore sequencing of *S. enterica*, a small amount of the microbial DNA standard (0.1 μl, 0.3 μl, 0.5 μl) was augmented with 2 μl of the lambda DNA standard to obtain approximately 600 ng of DNA (Table S3). The final concentrations of these noise-augmented samples meet the minimum DNA input requirement for Flongle flow cells (R9.4.1).

### Ligation of the augmented DNA samples

A ligation-based sequencing kit was chosen for processing singleplex samples of the noise-augmented target DNA. Library preparation was carried out using the ligation sequencing kits (SQK-LSK109; Oxford Nanopore Technologies) according to the manufacturer’s instructions. For Flongle flow cells (R9.4.1), the Flongle Sequencing Expansion (EXP-FSE001) was used in combination for optimal results. These ligation Sequencing kits are optimized for preparing sequencing libraries from dsDNA such as gDNA, cDNA, or amplicons. The library preparation method involves repairing and dA-tailing DNA ends using the NEBNext End Repair/dA-tailing module, and then ligating sequencing adapters onto the prepared ends. For the highest data yields, these ligation kits recommend starting with 1 μg of gDNA or 100-200 fmol of shorter-fragment input such as amplicons or cDNA. Starting with lower amounts of input material, or impure samples, may affect library preparation efficiency and reduce sequencing throughput.

### Nanopore sequencing of the NAD samples with adaptive sampling

The NAD samples of the microbial DNA standards augmented with the noise DNA were sequenced with a MinION Mk1B (Oxford Nanopore Technologies). For each sequencing run, adaptive sampling was preset to deplete the Lambda phage genome. The complete genome of *Escherichia coli* bacteriophage (NC_001416.1) was uploaded as a FASTA file as the reference sequence to deplete while sampling. Adaptive sampling requires high computational power due to the need to conduct real-time basecalling to decide whether to reject or accept reads for further sequencing. GPU-accelerated adaptive sampling was performed using an NVIDIA GPU on Windows (NVIDIA Quadro P3000).

### High-accuracy basecalling of NAD sequencing experiments

After the NAD sequencing of the samples was completed, the raw signal data in FAST5 files were basecalled with Guppy (v6.5.7). The GPU version of Guppy was used to improve the performance of super-accuracy basecalling, which achieves the highest raw read accuracy out of the other available neural-network models in Guppy, such as fast analysis and high-accuracy analysis [38]. An external GPU enclosure (eGPU) paired with an Nvidia Ampere card (RTX3060) was connected to a Dell Latitude laptop to perform high-accuracy basecalling, saving the basecalled long reads in FASTQ files.

### Rapid classification pipeline of NAD sequencing experiments

For the rapid classification of reads, a cloud-based platform providing analysis workflows called EPI2ME was used. Using the EPI2ME platform (v.3.5.7; Oxford Nanopore Technologies, Oxford, UK), the WIMP workflow (v.2021.11.26) rapidly classifies long reads from nanopore sequencing based on the Centrifuge classification engine [39]. The NAD datasets basecalled with the super-accuracy models were classified at the Family, Genus, and Species level using the WIMP workflow.

The classification of each long read was saved to analyze the performance of NAD sequencing in accuracy and efficiency. For accuracy, the decision of adaptive sampling to accept or reject further sequencing of each read was analyzed by the reads classified as the target (bacterial DNA) versus the noise (Lambda phage DNA). For efficiency, the mean DNA length sequenced for the reads classified as the target (bacterial DNA) versus the noise (Lambda phage DNA) was calculated.

### Genome assembly of the noise DNA and the target DNA

For the downstream analysis, the NAD datasets from each sample were assembled using Flye (v2.9.2) [40,41]. For the assembly, the FASTQ files generated from each experiment were combined into one file, and the metagenomic option was used to assemble the long reads into genomes [42].

After metagenomic genome assembly, the resulting contigs were aligned to the reference noise genome (NC_001416) and the reference target genome (GCF_000006945 or GCF_000006765) using MUMmer (v4.0+) [43–45]. The alignment quality of NAD datasets to a reference genome was analyzed and visualized using Assemblytics [46].

## Declarations

### Availability of data and materials

All codes related to this project are available under an open-source license at https://github.com/hshimlab. For data analysis, Python v.3.6.4 (https://www.python.org), NumPy v.1.17.5 (https://github.com/numpy/numpy), SciPy v.1.1.0 (https://www.scipy.org), seaborn v.0.9.0 (https://github.com/mwaskom/seaborn), Matplotlib v.3.3.4 (https://github.com/matplotlib/matplotlib), pandas v.0.22.0 (https://github.com/pandas-dev/pandas) were used. For nanopore data acquisition, we used the MinKNOW v.21.11.8 and MinKNOW core v.2.1.0. For rapid nanopore data analysis, we used the EPI2ME platform v.3.5.7. For high-accuracy basecalling, we used Guppy v6.5.7. For genome assembly and visualization, we used Flye v2.9.2, MUMmer v4.0+, and Assemblyticcs.

### Ethics approval and consent to participate

Not Applicable

### Consent for publication

Not Applicable

